# Deep immunological imprinting due to the ancestral spike in the current bivalent COVID-19 vaccine

**DOI:** 10.1101/2023.05.03.539268

**Authors:** Qian Wang, Yicheng Guo, Anthony R. Tam, Riccardo Valdez, Aubree Gordon, Lihong Liu, David D. Ho

## Abstract

With the aim of broadening immune responses against the evolving SARS-CoV-2 Omicron variants, bivalent COVID-19 mRNA vaccines that encode the ancestral and Omicron BA.5 spike proteins have been authorized for clinical use, supplanting the original monovalent counterpart in numerous countries. However, recent studies have demonstrated that administering either a monovalent or bivalent vaccine as a fourth vaccine dose results in similar neutralizing antibody titers against the latest Omicron subvariants, raising the possibility of immunological imprinting. Utilizing binding immunoassays, pseudotyped virus neutralization assays, and antigenic mapping, we investigated antibody responses from 72 participants who received three monovalent mRNA vaccine doses followed by either a bivalent or monovalent booster, or who experienced breakthrough infections with the BA.5 or BQ subvariant after vaccinations with an original monovalent vaccine. Compared to a monovalent booster, the bivalent booster did not yield noticeably higher binding titers to D614G, BA.5, and BQ.1.1 spike proteins, nor higher virus-neutralizing titers against SARS-CoV-2 variants including the predominant XBB.1.5 and the emergent XBB.1.16. However, sera from breakthrough infection cohorts neutralized Omicron subvariants significantly better. Multiple analyses of these results, including antigenic mapping, made clear that inclusion of the ancestral spike prevents the broadening of antibodies to the BA.5 component in the bivalent vaccine, thereby defeating its intended goal. Our findings suggest that the ancestral spike in the current bivalent COVID-19 vaccine is the cause of deep immunological imprinting. Its removal from future vaccine compositions is therefore strongly recommended.

## Main text

The FDA recently amended the emergency use authorization for the bivalent (ancestral/BA.5) COVID-19 mRNA vaccines to streamline the vaccination schedule and to allow older and immunocompromised individuals to receive additional booster shots^1^. However, several studies have reported that serum neutralizing antibody titers against SARS-CoV-2 Omicron BA.5 and subsequent subvariants after a bivalent vaccine booster were not discernibly better than after a monovalent (ancestral) booster^2-4^. We now present new findings and analyses to show that the ancestral spike exacerbates immunological imprinting and should be eliminated from future vaccine compositions.

We collected serum from 72 individuals who had received three doses of vaccines followed by a monovalent or bivalent booster or who had experienced a BA.5 or BQ breakthrough infection. Clinical details for all cases are provided in **Table S1** and summarized in **Table S2**. Each serum sample was tested in pseudovirus assays to determine neutralizing antibody titers against the ancestral D614G strain and a panel of Omicron subvariants, including BA.2, BA.5, BQ.1.1, CH.1.1, XBB.1.5, and the newly surging XBB.1.16. We also performed immunoassays to quantify serum antibodies that bind the spike proteins of D614G, BA.5, and BQ.1.1.

Each cohort exhibited roughly similar (<2-fold difference) serum binding antibody titers to D614G, BA.5, and BQ.1.1 spike proteins (**Figure S1**). As for serum SARS-CoV-2-neutralizing antibodies, all cohorts had the highest titers against D614G but substantially lower titers against the Omicron subvariants, particularly the currently dominant XBB.1.5 and the emergent XBB.1.16 (**Figure S2**). Notably, the extent of antibody evasion exhibited by XBB.1.16 and its spike point mutants (E180V and K478R) was comparable to that of XBB.1.5 (**Figure S3**). The data in **Figure S2** were further analyzed in three ways. First, antigenic cartography revealed that sera from the “monovalent” and “bivalent” cohorts were substantially overlapping and centered around the ancestral strain, whereas sera from BA.5 and BQ breakthrough cohorts were similarly shifted toward BA.5 and BQ.1.1 (**Figure 1A**). Second, the above findings prompted replotting of a subset of the data to specifically examine the issue of immunological imprinting^5^ (**Figure 1B**). Serum neutralizing antibodies against D614G were similar for all cohorts, with small differences that did not reach statistical significance. However, titers against BA.5 or BQ.1.1 were significantly higher for BA.5 (3.4 to 3.7-fold) and BQ.1.1 (3.6 to 4.5-fold) breakthrough cohorts. These findings made clear the role of the ancestral spike in immunological imprinting in that exposure of the immune system to both the ancestral and BA.5 spikes did not elicit discernibly better BA.5-neutralizing antibodies, whereas exposure to only BA.5 spike (through infection) in the absence of the ancestral spike did elicit such antibodies. That BQ1.1 and BA.5 results were similar should not be surprising since BQ.1.1 is a direct descendant of BA.5. Third, we created antigenic maps based on the neutralization data for each of the clinical cohorts (**Figure 1C**). The antigenic distances from D614G to BA.5 or to BQ.1.1 were similar for the “monovalent” and “bivalent” vaccine cohorts. In contrast, these antigenic distances were substantially shortened with BA.5 or BQ breakthrough infection, and these differences reached high-level statistical significance (**Figure S4**). This analysis graphically demonstrates that inclusion of the ancestral spike in the bivalent vaccine precludes the broadening of neutralizing antibodies to BA.5, which was clearly evident in the breakthrough infection cases.

**Figure 1.**
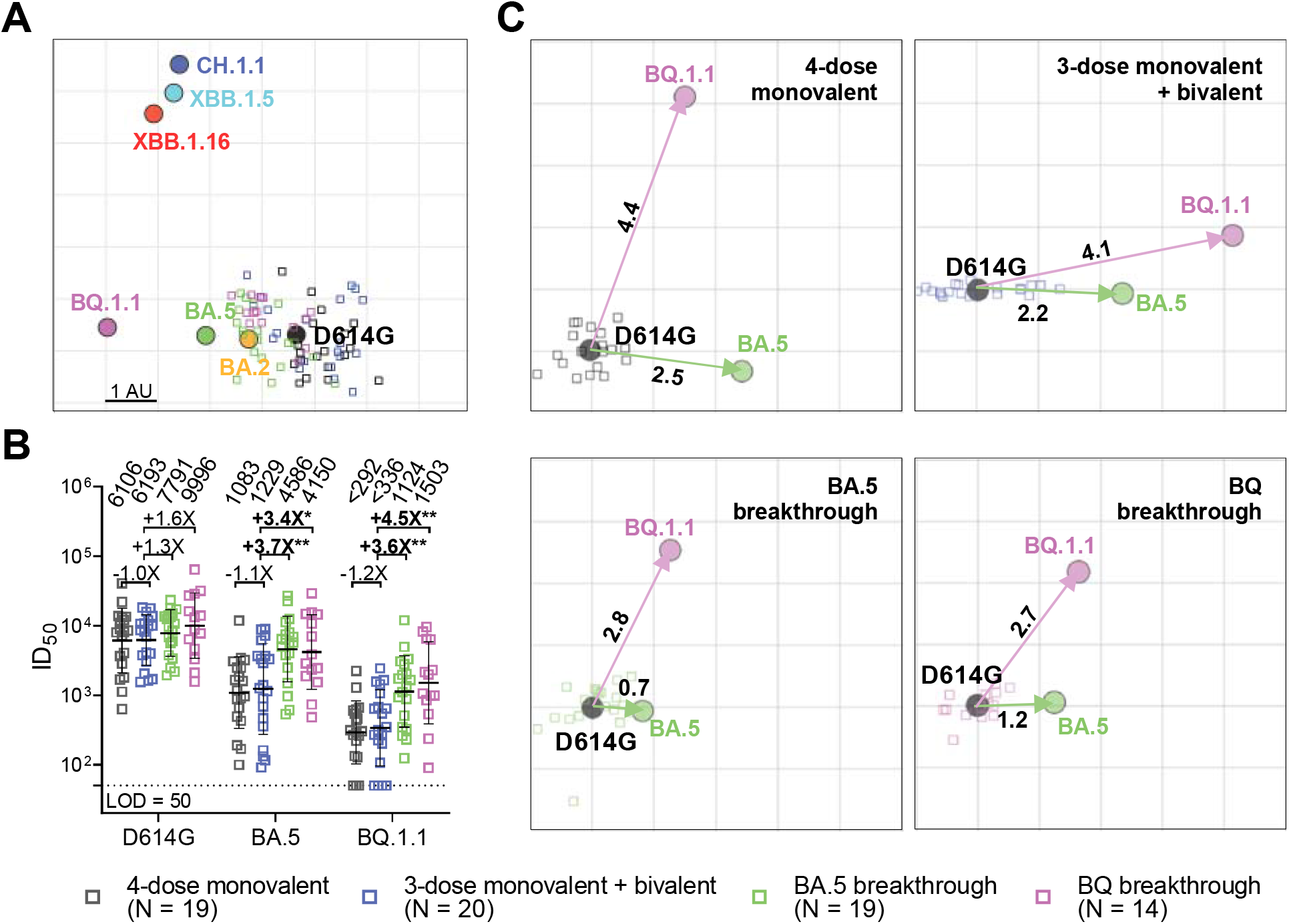
SARS-CoV-2 neutralizing antibody responses following monovalent booster, bivalent booster, or breakthrough infection. Panel **A** presents antigenic map derived from the neutralization data of the serum samples from participants who received four doses of a monovalent mRNA vaccine (4-dose monovalent), three doses of a monovalent mRNA vaccine followed by one dose of a bivalent vaccine targeting the ancestral virus and BA.5 (3-dose monovalent + bivalent), and who experienced either BA.5 (BA.5 breakthrough) or BQ (BQ breakthrough) breakthrough infections after two to four doses of vaccine. SARS-CoV-2 variants and sera are shown as colored circles and squares, respectively. The X and Y axes represent antigenic units (AU), with each grid corresponding to a two-fold serum dilution of the neutralization titer. Panel **B** displays the neutralizing antibody responses induced by a fourth dose of a bivalent mRNA vaccine compared to a fourth dose of the original monovalent booster or breakthrough infections. The values above the symbols indicate the geometric mean ID_50_ titer (GMT) for each cohort. The assay limit of detection (LOD = 50) is represented by a dotted line. Mann-Whitney test was used to compare the results and the fold-changes in GMT are also shown. The statistical significance of the results is represented as \**p* < 0.05 or \*\**p* < 0.01. Panel **C** displays the antigenic maps for individual cohorts against D614G, BA.5, and BQ.1.1. Arrows indicate the antigenic distances from D614G to BA.5 (green) and BQ.1.1 (magenta).

Much of the world’s population has been immunologically exposed to the ancestral spike of SARS-CoV-2 through either vaccination or infection, or both. The inclusion of this spike in our COVID-19 vaccines will continue to skew our antibody responses toward what we have already seen and away from what we wish to elicit going forward. Therefore, based on observations made herein, we put forth a strong recommendation to remove the ancestral spike from new COVID-19 vaccines for the foreseeable future.

## References

1. Coronavirus (COVID-19) Update: FDA Authorizes Changes to Simplify Use of Bivalent mRNA COVID-19 Vaccines. U.S. Food & Drug Administration, 2023. at https://www.fda.gov/news-events/press-announcements/coronavirus-covid-19-update-fda-authorizes-changes-simplify-use-bivalent-mrna-covid-19-vaccines.)

2. Wang Q, Bowen A, Valdez R, et al. Antibody Response to Omicron BA.4-BA.5 Bivalent Booster. N Engl J Med 2023;388:567–9.

3. Wang Q, Bowen A, Tam AR, et al. SARS-CoV-2 neutralising antibodies after bivalent versus monovalent booster. Lancet Infect Dis 2023;23:527–8.

4. Collier AY, Miller J, Hachmann NP, et al. Immunogenicity of BA.5 Bivalent mRNA Vaccine Boosters. N Engl J Med 2023;388:565–7.

5. Wheatley AK, Fox A, Tan HX, et al. Immune imprinting and SARS-CoV-2 vaccine design. Trends Immunol 2021;42:956–9.

