## Supplemental information for "Deep immunological imprinting due to the ancestral spike in the current bivalent COVID-19 vaccine"

### Supplementary Appendix

#### Supplementary Methods

*Clinical cohorts*

The sera analyzed in this study were obtained from four cohorts: the "4-dose monovalent", "3-dose monovalent + bivalent", "BA.5 breakthrough" and "BQ breakthrough". The first cohort consisted of individuals who received four doses of the monovalent COVID-19 mRNA vaccine, whereas the second cohort received three doses of the monovalent COVID-19 mRNA vaccine followed by one dose of the Pfizer or Moderna bivalent mRNA vaccine. The last two cohorts comprised of patients who had Omicron BA.5 or BQ breakthrough infection after vaccination. To determine prior SARS-CoV-2 infection status, serum samples were evaluated using anti-nucleocapsid protein (NP) ELISA.

The study collected sera from two sources. The first source of sera was obtained from the Immunity-Associated with SARS-CoV-2 Study (IASO), an ongoing cohort study that began in 2020 at the University of Michigan[1]. The participants in the IASO study provided written informed consent, and the serum samples were collected under a protocol approved by the Institutional Review Board of the University of Michigan Medical School. The second source was Columbia University Irving Medical Center, where a subset of vaccinee and breakthrough sera were collected. All subjects provided written informed consent, and the serum collections were performed under protocols reviewed and approved by the Institutional Review Board of Columbia University. More detailed information for each case can be found in **Table S1**, while a summary is provided in **Table S2**.

*Cell lines*

The Vero-E6 cells (CRL-1586) and HEK293T cells (CRL-3216) were obtained from the American Type Culture Collection (ATCC). The cells were cultured in Dulbecco's Modified Eagle Medium (DMEM) supplemented with 10% fetal bovine serum and 1% penicillin-streptomycin. The cells were maintained in a 5% $CO_2$ atmosphere at 37°C.

*SARS-CoV-2 spike plasmids*

Plasmids that encode the spike proteins of various SARS-CoV-2 variants, including D614G, BA.2, BA.5, BQ.1.1, and XBB.1.5, were previously created[2-6]. The spikes XBB.1.16, XBB.1.5-E180V, and XBB.1.5-K478R were generated using the QuikChange II XL site-directed mutagenesis kit according to the manufacturer's instructions (Agilent). Before use in experiments, the sequences of all constructs were verified by Sanger sequencing.

*Pseudovirus production*

The pseudotyped SARS-CoV-2 variants were created by replacing the native glycoprotein of the vesicular stomatitis virus (VSV) with the spike protein of the relevant SARS-CoV-2 variant[7]. To achieve this, HEK293T cells were transfected with a plasmid that encodes the relevant spike protein using polyethyleneimine (PEI) at a concentration of 1 mg/mL. The transfected cells were then incubated at 37°C in a 5% $CO_2$ atmosphere for 24 hours, followed by infection with the VSV-G pseudotyped ΔG-luciferase (G*ΔG-luciferase, Kerafast). Following a 2-hour incubation period at 37°C, the infected HEK293T cells were washed three times and cultured in fresh medium for an additional 24 hours. The resulting supernatants were collected, centrifuged to remove any precipitates, and stored at -80°C. To prevent any contamination from VSV-G pseudotyped ΔG-luciferase, the viral stock was pre-incubated with 20% I1 hybridoma (anti-VSV-G) supernatant (ATCC; CRL-2700) for 1 hour at 37°C before infecting the target cells.

*Pseudovirus neutralization*

To ensure consistent viral input, the pseudovirus titers were standardized before conducting the neutralization assay. The heat-inactivated sera were then tested in triplicate using 96-well plates and were serially diluted four-fold in media, beginning with a 1:50 dilution. After the sera were diluted, they were combined with the pseudoviruses and incubated at 37°C for 1 hour. Vero-E6 cells were then added to each well at a density of $3 \times 10^4$ cells/well and were incubated at 37°C with 5% $CO_2$ for 10 hours. Following the incubation, the cells were lysed, and luciferase activity was measured using the Luciferase Assay System (Promega) and SoftMax Pro v.7.0.2 (Molecular Devices), according to the manufacturer's instructions.

*ELISA*

The S2P spike trimer protein of D614G, BA.5, and BQ.1.1 were generated according to previously described methods[2]. To determine the binding titers of serum samples to D614G, BA.5, and BQ.1.1 spikes, we immobilized 50 ng/well of S2P trimer onto ELISA plates and incubated them overnight at 4°C. Subsequently, the ELISA plates were blocked with 300 µl blocking buffer (1% BSA and 10% bovine calf serum (BCS) (Sigma)) in PBS at 37°C for 2 hours. Afterwards, serum samples were serially diluted by 5-fold from 100× using dilution buffer (1% BSA and 20% BCS in PBS) and incubated in the ELISA plates at 37°C for 1 h. After incubation, 10,000-fold diluted Peroxidase AffiniPure goat anti-human IgG (H+L) antibody (Jackson ImmunoResearch) was added and incubated for 1 h at 37°C. The plates were washed between each step with PBST (0.5% Tween-20 in PBS). Finally, the TMB substrate (Sigma) was added and incubated before the reaction was stopped using 1 M sulfuric acid. Absorbance was measured at 450 nm. $EC_{50}$ values was calculated as the dilutions at which the OD450 readings reached half of the maximal using GraphPad Prism v.9.2.

*Antigenic cartography*

Antigenic distances between SARS-CoV-2 variants were estimated by integrating the neutralization potency of each serum sample using a previously described antigenic cartography method[8]. The map was generated with the Racmacs package (https://acorg.github.io/Racmacs/, v.1.1.4) in R, using 2,000 optimization steps and setting the minimum column basis parameter to 'none'. The mapDistances function in the Racmacs package was employed to calculate antigenic distances, and the average distances for all sera to variants were used to represent the final distances. D614G served as the center of sera for each group, the seeds for each antigenic map were manually adjusted to ensure that BA.5 was displayed in the horizontal direction relative to the sera.

#### Acknowledgements

We thank Sho Iketani for providing BQ.1.1 S2P spike trimer protein. This study was supported financially by the SARS-CoV-2 Assessment of Viral Evolution Program, NIAID, NIH (Subcontract No. 0258-A709-4609 under Federal Contract No. 75N93021C00014) awarded to D.D.H., as well as the NIAID, NIH (Subcontract under Contract Number 75N93019C00051) awarded to A.G. The authors express their gratitude to David Manthei, Carmen Gherasim, Emily Stoneman, Adam Lauring, Victoria Blanc, Pamela Bennett-Baker, Savanna Sneeringer, Theresa Kowalski-Dobson, Alyssa Meyers, Zijin Chu, Hailey Kuiken, Lonnie Barnes, Ashley Eckard, Kathleen Lindsey, Dawson Davis, Aaron Rico, Daniel Raymond, Mayurika Patel, and Nivea Vydiswaran from the IASO study team for their contribution in providing the serum samples.

#### Author Contributions

L.L. and D.D.H. conceived the study. Q.W., A.R.T., and L.L. performed experiments. R.V. and A.G. collected serum samples. Y.G. generated antigenic cartography. Q.W., Y.G., L.L., and D.D.H. analyzed the results and wrote the manuscript. L.L. and D.D.H. directed and supervised the project. All authors reviewed and approved of the manuscript.

#### Declaration of Interests

The authors declare potential conflicts of interest as follows: D.D.H. is a co-founder of TaiMed Biologics and RenBio, as well as a board director for Vicarious Surgical; he also serves as a consultant to WuXi Biologics, Brii Biosciences, and Veru; and he receives research support from Regeneron. A.G. served on a scientific advisory board for Janssen Pharmaceuticals. The remaining authors declare no competing interests.

**Table S1. Demographics of clinical cohorts**

| Sample ID | Vaccine type and infected strain | Days post-vaccination or *infection | Confirmed COVID-19 | Age | Gender |
|---|---|---|---|---|---|
| *4-dose monovalent* | | | | | |
| UM-65 | BNT162b2/BNT162b2/BNT162b2/BNT162b2 | 24 | No | 52 | Female |
| UM-66 | BNT162b2/BNT162b2/BNT162b2/BNT162b2 | 20 | No | 57 | Female |
| UM-67 | BNT162b2/BNT162b2/BNT162b2/BNT162b2 | 20 | No | 61 | Female |
| UM-68 | mRNA-1273/mRNA-1273/mRNA-1273/mRNA-1273 | 22 | No | 48 | Female |
| UM-69 | BNT162b2/BNT162b2/BNT162b2/BNT162b2 | 23 | No | 50 | Female |
| UM-70 | BNT162b2/BNT162b2/BNT162b2/BNT162b2 | 22 | No | 50 | Female |
| UM-71 | BNT162b2/BNT162b2/BNT162b2/BNT162b2 | 20 | No | 58 | Female |
| UM-72 | BNT162b2/BNT162b2/BNT162b2/BNT162b2 | 26 | No | 56 | Female |
| UM-73 | BNT162b2/BNT162b2/BNT162b2/BNT162b2 | 29 | No | 63 | Female |
| UM-74 | BNT162b2/BNT162b2/BNT162b2/BNT162b2 | 25 | No | 58 | Female |
| UM-75 | BNT162b2/BNT162b2/BNT162b2/BNT162b2 | 21 | No | 62 | Male |
| UM-76 | BNT162b2/BNT162b2/BNT162b2/BNT162b2 | 26 | No | 54 | Female |
| UM-77 | BNT162b2/BNT162b2/BNT162b2/BNT162b2 | 23 | No | 53 | Male |
| UM-78 | BNT162b2/BNT162b2/BNT162b2/BNT162b2 | 21 | No | 55 | Female |
| UM-79 | BNT162b2/BNT162b2/BNT162b2/BNT162b2 | 23 | No | 59 | Female |
| UM-80 | BNT162b2/BNT162b2/BNT162b2/BNT162b2 | 21 | No | 49 | Female |
| UM-81 | BNT162b2/BNT162b2/BNT162b2/BNT162b2 | 27 | No | 57 | Female |
| UM-82 | BNT162b2/BNT162b2/BNT162b2/BNT162b2 | 27 | No | 55 | Female |
| Q97 | BNT162b2/BNT162b2/BNT162b2/BNT162b2 | 36 | No | 53 | Female |
| *3-dose monovalent + bivalent* | | | | | |
| UM-36 | BNT162b2/BNT162b2/BNT162b2/Moderna Bivalent | 24 | No | 38 | Female |
| UM-37 | BNT162b2/BNT162b2/BNT162b2/Moderna Bivalent | 27 | No | 42 | Female |
| UM-39 | mRNA-1273//mRNA-1273/mRNA-1273/Moderna Bivalent | 24 | No | 36 | Male |
| UM-40 | BNT162b2/BNT162b2/BNT162b2/Pfizer Bivalent | 25 | No | 37 | Female |
| UM-43 | BNT162b2/BNT162b2/BNT162b2/Pfizer Bivalent | 25 | No | 49 | Female |
| UM-44 | BNT162b2/BNT162b2/BNT162b2/Moderna Bivalent | 25 | No | 37 | Female |
| UM-47 | BNT162b2/BNT162b2/BNT162b2/Pfizer Bivalent | 26 | No | 45 | Male |
| UM-48 | BNT162b2/BNT162b2/mRNA-1273/Moderna Bivalent | 26 | No | 43 | Female |
| UM-51 | mRNA-1273/mRNA-1273/mRNA-1273/Moderna Bivalent | 29 | No | 32 | Female |
| UM-52 | BNT162b2/BNT162b2/BNT162b2/Pfizer Bivalent | 23 | No | 43 | Female |
| UM-53 | BNT162b2/BNT162b2/BNT162b2/Pfizer Bivalent | 26 | No | 43 | Female |
| UM-54 | BNT162b2/BNT162b2/mRNA-1273/Moderna Bivalent | 29 | No | 38 | Female |
| UM-55 | BNT162b2/BNT162b2/BNT162b2/Moderna Bivalent | 28 | No | 38 | Female |
| UM-56 | BNT162b2/BNT162b2/mRNA-1273/Moderna Bivalent | 27 | No | 36 | Female |
| UM-60 | BNT162b2/BNT162b2/BNT162b2/Moderna Bivalent | 30 | No | 24 | Female |
| Q101 | mRNA-1273/mRNA-1273/mRNA-1273/Moderna Bivalent | 30 | No | 32 | Female |
| Q102 | BNT162b2/BNT162b2/mRNA-1273/Moderna Bivalent | 23 | No | 39 | Male |
| Q103 | BNT162b2/BNT162b2/BNT162b2/Pfizer Bivalent | 30 | No | 26 | Female |
| Q104 | mRNA-1273/mRNA-1273/mRNA-1273/Pfizer Bivalent | 30 | No | 27 | Female |
| Q105 | BNT162b2/BNT162b2/BNT162b2/Pfizer Bivalent | 23 | No | 23 | Male |
| *BA.5 breakthrough* | | | | | |
| Q71 | mRNA-1273/mRNA-1273/BNT162b2/BA.5.2.1 | *29 | Yes | 29 | Female |
| Q77 | BNT162b2/BNT162b2/BNT162b2/BA.5 | *22 | Yes | 61 | Female |
| Q79 | mRNA-1273/mRNA-1273/mRNA-1273/BA.5 | *15 | Yes | 28 | Female |
| Q80 | mRNA-1273/mRNA-1273/mRNA-1273/BA.5 | *21 | Yes | 24 | Female |
| Q81 | BNT162b2/BNT162b2/BNT162b2/BA.5 | *75 | Yes | 35 | Female |
| Q82 | BNT162b2/BNT162b2/mRNA-1273/BA.5 | *63 | Yes | 46 | Female |
| Q83 | BNT162b2/BNT162b2/BNT162b2/BA.5 | *28 | Yes | 55 | Male |
| Q84 | BNT162b2/BNT162b2/BNT162b2/BA.5 | *17 | Yes | 57 | Female |
| UM-85 | BNT162b2/BNT162b2/BNT162b2/BA.5 | *29 | Yes | 44 | Female |

| Sample ID | Vaccine type and infected strain | Days post-vaccination or *infection | Confirmed COVID-19 | Age | Gender |
|---|---|---|---|---|---|
| UM-86 | BNT162b2/BNT162b2/mRNA-1273/BA.5 | *29 | Yes | 36 | Female |
| UM-87 | BNT162b2/BNT162b2/BNT162b2/BNT162b2/BA.5 | *31 | Yes | 54 | Female |
| UM-88 | BNT162b2/BNT162b2/BNT162b2/BNT162b2/BA.5 | *28 | Yes | 69 | Male |
| UM-89 | BNT162b2/BNT162b2/BNT162b2/BNT162b2/BA.5 | *42 | Yes | 44 | Male |
| UM-90 | BNT162b2/BNT162b2/BNT162b2/BNT162b2/BA.5 | *28 | Yes | 41 | Female |
| UM-91 | BNT162b2/BNT162b2/BNT162b2/BNT162b2/BA.5 | *28 | Yes | 44 | Female |
| UM-92 | BNT162b2/BNT162b2/BNT162b2/BNT162b2/BA.5 | *31 | Yes | 29 | Female |
| UM-93 | BNT162b2/BNT162b2/BNT162b2/BNT162b2/BA.5 | *29 | Yes | 48 | Female |
| UM-96 | BNT162b2/BNT162b2/BNT162b2/BNT162b2/BA.5 | *33 | Yes | 58 | Female |
| Q106 | mRNA-1273/mRNA-1273/mRNA-1273/mRNA-1273/BA.5 | *18 | Yes | 32 | Female |
| *BQ breakthrough* | | | | | |
| BQ-1 | BNT162b2/BNT162b2/BNT162b2/BNT162b2/mRNA-1273/BQ | *33 | Yes | 53 | Female |
| BQ-2 | BNT162b2/BNT162b2/BNT162b2/mRNA-1273/BQ | *30 | Yes | 32 | Female |
| BQ-3 | BNT162b2/BNT162b2/BNT162b2/mRNA-1273/BQ | *21 | Yes | 35 | Female |
| BQ-4 | BNT162b2/BNT162b2/mRNA-1273/BQ | *39 | Yes | 52 | Female |
| BQ-5 | BNT162b2/BNT162b2/BNT162b2/BNT162b2/mRNA-1273/BQ | *22 | Yes | 62 | Female |
| BQ-6 | BNT162b2/BNT162b2/mRNA-1273/BQ | *34 | Yes | 29 | Female |
| BQ-7 | BNT162b2/BNT162b2/BQ | *44 | Yes | 35 | Female |
| BQ-8 | mRNA-1273/mRNA-1273/mRNA-1273/mRNA-1273/BQ | *59 | Yes | 33 | Female |
| BQ-9 | BNT162b2/BNT162b2/BNT162b2/BNT162b2/BQ | *44 | Yes | 45 | Female |
| BQ-10 | JNJ-78436735/JNJ-78436735/BNT162b2/BNT162b2/BQ | *32 | Yes | 47 | Female |
| BQ-11 | BNT162b2/BNT162b2/BNT162b2/BQ | *36 | Yes | 36 | Female |
| BQ-12 | BNT162b2/BNT162b2/BNT162b2/BNT162b2/BNT162b2/BQ | *77 | Yes | 59 | Female |
| BQ-13 | BNT162b2/BNT162b2/BNT162b2/BNT162b2/BQ | *25 | Yes | 39 | Female |
| BQ-14 | BNT162b2/BNT162b2/BNT162b2/BNT162b2/BQ | *67 | Yes | 43 | Female |

**Table S2. Summary of clinical cohorts**

| Characteristic | 4-dose Monovalent (N=19) | 3-dose monovalent + bivalent (N=20) | BA.5 Breakthrough (N=19) | BQ Breakthrough (N=14) |
|---|---|---|---|---|
| Sex - No. (%) | | | | |
| Female | 17 (89.5%) | 16 (80.0%) | 16 (84.2%) | 14 (100%) |
| Male | 2 (10.5%) | 4 (20.0%) | 3 (15.8%) | 0 (0%) |
| Mean Age (range)-year | 55.3 (48 - 63) | 36.4 (23 - 49) | 43.9 (24 - 69) | 42.9 (29 - 62) |
| Days post-vaccination or *infection | 24.0 | 26.5 | *31.4 | *40.2 |

*For the BA.5 breakthrough cohort, days are measured between last dose of vaccine and documented breakthrough infection.

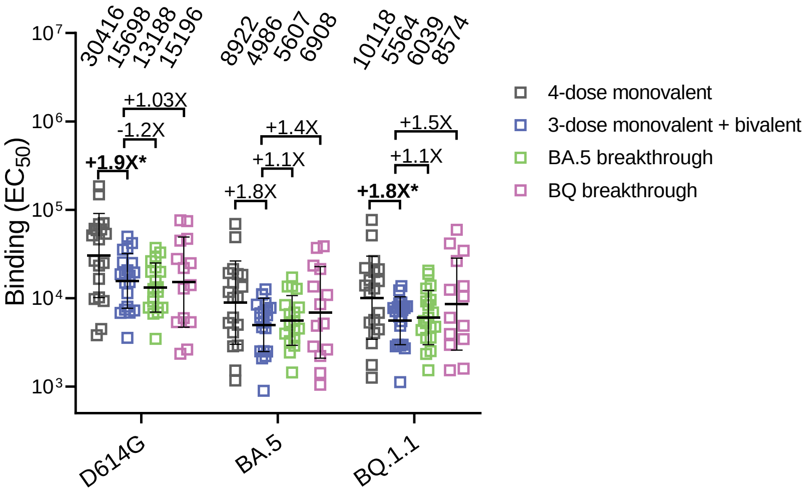

**Figure S1. SARS-CoV-2 binding antibody responses to D614G, BA.5, and BQ.1.1 spikes following monovalent booster, bivalent booster, or breakthrough infection.**

$EC_{50}$ titers of binding antibodies in the serum samples from participants form "4-dose monovalent", "4-dose monovalent + bivalent", "BA.5 breakthrough" and "BQ breakthrough" cohorts. The values above the symbols indicate the geometric mean $EC_{50}$ titers for each cohort. Mann-Whitney test was used to compare the results for the "3-dose monovalent + bivalent cohort" and the fold changes in geometric mean $EC_{50}$ titers are also shown. *$p < 0.05$.

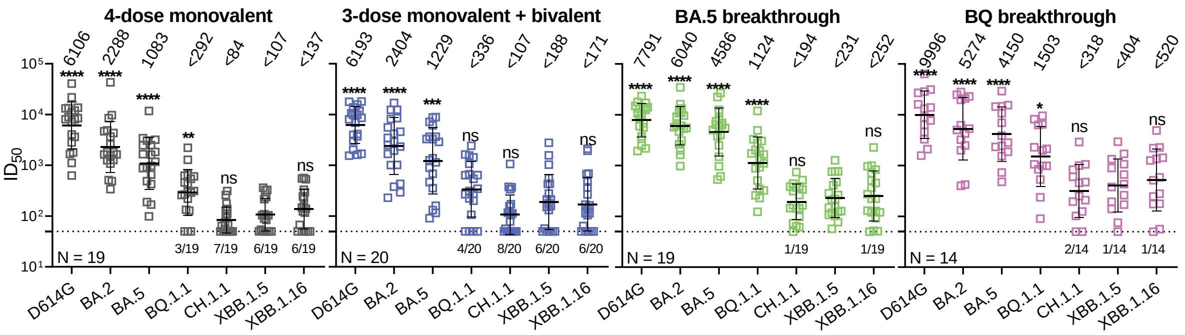

**Figure S2. SARS-CoV-2 neutralizing antibody responses following monovalent booster, bivalent booster, or breakthrough infection.**

$ID_{50}$ titers of neutralizing antibodies in the serum samples from participants who received four doses of a monovalent mRNA vaccine (4-dose monovalent), three doses of a monovalent mRNA vaccine followed by one dose of a bivalent vaccine targeting BA.5 variants (4-dose monovalent + bivalent), and experienced BA.5 (BA.5 breakthrough) or BQ (BQ breakthrough) breakthrough infections after two to four doses of vaccine. The values above the symbols indicate the geometric mean $ID_{50}$ titer (GMT) for each cohort. The assay limit of detection (LOD = 50) is represented by a dotted line, and the number of samples at or below the LOD is indicated above the x-axis. Mann-Whitney test was used to compare the results with XBB.1.5. ns, not significant; *$p < 0.05$; **$p < 0.01$; ***$p < 0.001$; ****$p < 0.0001$.

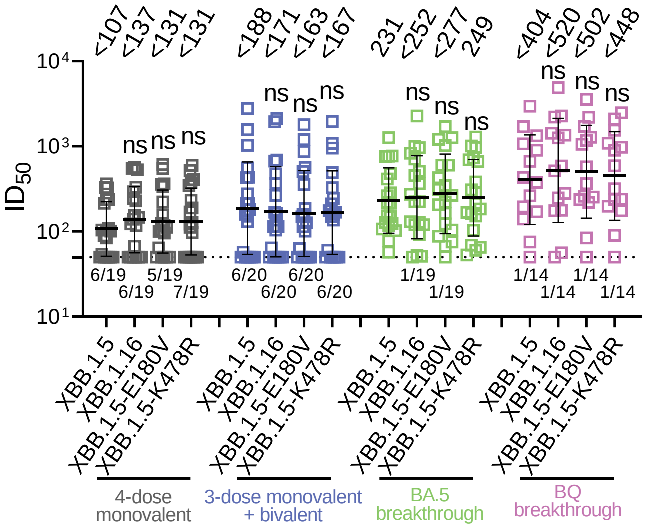

**Figure S3. Serum neutralizing ID50 titers against pseudotyped viruses of XBB.1.5, XBB.1.16, and XBB.1.5 carrying the individual spike mutations found in XBB.1.16.**

Values above symbols denote the geometric mean ID50 titer. The assay limit of detection (LOD) is 50 (dotted line) and the number of samples below the LOD is denoted above the x-axis. Comparisons were made against XBB.1.5 by Mann-Whitney tests. ns, not significant.

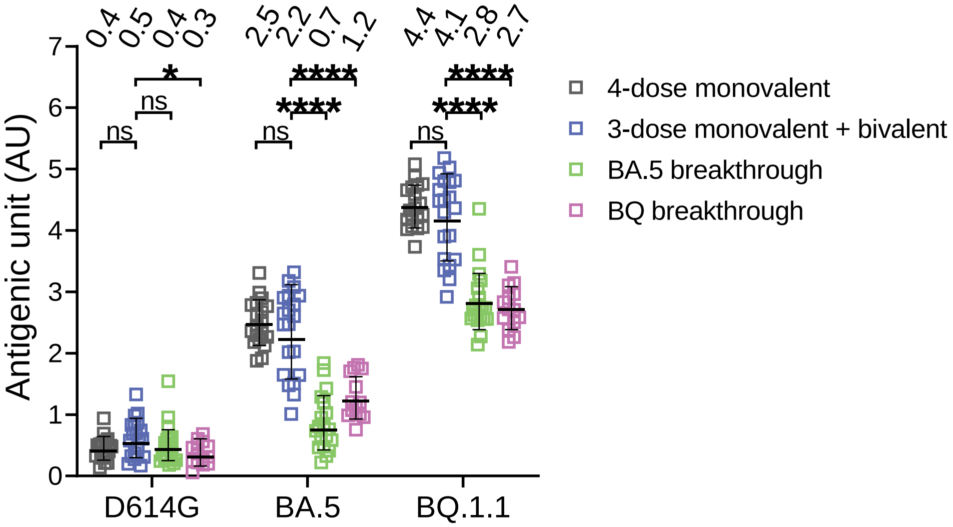

**Figure S4. Antigenic units from D614G, BA.5, and BQ.1.1 to the serum samples in the "4-dose monovalent", "4-dose monovalent + bivalent", "BA.5 breakthrough" and "BQ breakthrough" cohorts.**

Data were generated based on the neutralization data from **Figure S1**. One AU corresponds to a two-fold serum dilution of the neutralization titer. Mann-Whitney test was used to compare the results for the "3-dose monovalent + bivalent cohort". ns, not significant; $*p < 0.05$; $****p < 0.0001$.

#### Supplementary References

1. Simon V, Kota V, Bloomquist RF, et al. PARIS and SPARTA: Finding the Achilles' Heel of SARS-CoV-2. *mSphere* 2022; **7**(3): e0017922.
2. Wang Q, Iketani S, Li Z, et al. Alarming antibody evasion properties of rising SARS-CoV-2 BQ and XBB subvariants. *Cell* 2023; **186**(2): 279-86 e8.
3. Liu L, Iketani S, Guo Y, et al. Striking antibody evasion manifested by the Omicron variant of SARS-CoV-2. *Nature* 2022; **602**(7898): 676-81.
4. Iketani S, Liu L, Guo Y, et al. Antibody evasion properties of SARS-CoV-2 Omicron sublineages. *Nature* 2022; **604**(7906): 553-6.
5. Wang Q, Guo Y, Iketani S, et al. Antibody evasion by SARS-CoV-2 Omicron subvariants BA.2.12.1, BA.4, & BA.5. *Nature* 2022.
6. Wang Q, Bowen A, Tam AR, et al. SARS-CoV-2 neutralising antibodies after bivalent versus monovalent booster. *Lancet Infect Dis* 2023; **23**(5): 527-8.
7. Liu L, Wang P, Nair MS, et al. Potent neutralizing antibodies against multiple epitopes on SARS-CoV-2 spike. *Nature* 2020; **584**(7821): 450-6.
8. Smith DJ, Lapedes AS, de Jong JC, et al. Mapping the antigenic and genetic evolution of influenza virus. *Science* 2004; **305**(5682): 371-6.
